# Logistic proliferation of cells in scratch assays is delayed

**DOI:** 10.1101/077388

**Authors:** Wang Jin, Esha T Shah, Catherine J Penington, Scott W McCue, Philip K Maini, Matthew J Simpson

## Abstract

Scratch assays are used to study how a population of cells recolonises a vacant region on a two-dimensional substrate after a cell monolayer is scratched. These experiments are used in many applications including drug design for the treatment of cancer and chronic wounds. To provide insights into the mechanisms that drive scratch assays, solutions of continuum reaction–diffusion models have been calibrated to data from scratch assays. These models typically include a logistic source term to describe carrying capacity-limited proliferation, however the choice of using a logistic source term is often made without examining whether it is valid. Here we study the proliferation of PC-3 prostate cancer cells in a scratch assay. All experimental results for the scratch assay are compared with equivalent results from a proliferation assay where the cell monolayer is not scratched. Visual inspection of the time evolution of the cell density away from the location of the scratch reveals a series of sigmoid curves that could be naively calibrated to the solution of the logistic growth model. However, careful analysis of the per capita growth rate as a function of density reveals several key differences between the proliferation of cells in scratch and proliferation assays. Our findings suggest that the logistic growth model is valid for the entire duration of the proliferation assay. On the other hand, guided by data, we suggest that there are two phases of proliferation in a scratch assay; at short time we have a *disturbance phase* where proliferation is not logistic, and this is followed by a *growth phase* where proliferation appears to be logistic. These two phases are observed across a large number of experiments performed at different initial cell densities. Overall our study shows that simply calibrating the solution of a continuum model to a scratch assay might produce misleading parameter estimates, and this issue can be resolved by making a distinction between the disturbance and growth phases. Repeating our procedure for other scratch assays will provide insight into the roles of the disturbance and growth phases for different cell lines and scratch assays performed on different substrates.

## 1 Introduction

Understanding population dynamics is a fundamental question that has wide relevance to many biological and ecological processes. For example, the rate of spatial spreading of invasive species through different ecosystems is driven, in part, by the population dynamics and rates of growth of the invasive species (Lewis and Kareiva, 1993; Murray, 2002; Waters et al. 2015). Population dynamics and population growth are also central to understanding the spread of infectious diseases. For example, the spread of *Wolbachia* into wild mosquito populations is thought to reduce a wide range of diseases, and the spatial spreading of the mosquito population is partly driven by the population dynamics of the mosquito population (Chan and Kim, 2013). Similar ideas also apply to the spreading of tumour cells and the progression of cancer, which is related to the rates of proliferation of invasive cancer cells (Alarócn et al. 2003; Mallet and de Pillis 2006; Ribba et al. 2006). Therefore, improving our understanding of population dynamics by calibrating mathematical models to experimental observations of population dynamics is of great interest.

In *vitro* scratch assays are routinely used to study the ability of cell populations to re–colonise an initially–vacant region (Liang et al., 2007; Tremel et al., 2009; Kramer et al., 2013; Treloar and Simpson, 2013). This re–colonisation occurs as a result of the combination of cell migration and cell proliferation, and gives rise to moving fronts of cells that re–colonise the vacant region. Scratch assays provide insights into both cancer spreading and tissue repair processes (Maini et al., 2004a; Maini et al., 2004b; Kramer et al., 2013). In general, performing a scratch assay involves three steps: (i) growing a monolayer of cells on a two–dimensional substrate; (ii) creating a vacant region in the monolayer by scratching it with a sharp–tipped instrument; and, (iii) imaging the re–colonisation of the scratched region (Liang et al., 2007; Kramer et al., 2013). Another type of *in vitro* assay, called a proliferation assay, is performed using the exact same procedure as a scratch assay, except that the monolayer of cells is not scratched (Jones et al., 2001; Tremel et al., 2009; Simpson et al., 2013). Cell proliferation assays allow experimentalists to measure the increase in cell numbers over time due to proliferation (Tremel et al., 2009).

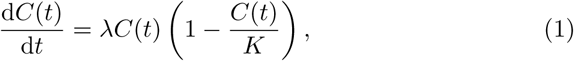

where *C*(*t*) > 0 is the density of cells, *t* is time, λ > 0 is the proliferation rate, and *K* > 0 is the carrying capacity density.

It is interesting to note that a logistic growth term is often used when modelling scratch assays or proliferation assays (Maini et al., 2004a; Maini et al., 2004b; Savla et al., 2004; Cai et al., 2007; Sengers et al., 2007; Tremel et al., 2009; Johnston et al., 2015; Jin et al., 2016), yet the suitability of this choice is rarely, if ever, tested using experimental data. In fact, several studies argue that the logistic growth equation does not always match experimental data (Laird, 1964;Zwietering et al., 1990;West et al., 2001;Sarapata and de Pillis, 2014). For example, Laird examines *in vivo* tumour growth data and shows that the standard logistic model does not match experimental data (Laird, 1964). Similarly,Sarapata and de Pillis find that the logistic growth model does not always match experimental tumour growth data (Sarapata and de Pillis, 2014).West and coworkers investigate the growth patterns of a wide range of animal models (West et al., 2001). By comparing experimental data with model predictions, they suggest that the growth is not logistic, and is beter described by a more general model. In addition, the results from our previous study, focusing on scratch assays, suggest that when calibrating solutions of a logistic–type reaction–diffusion equation to experimental data with varying initial cell density, there appears to be no unique value of λ for which the logistic growth equation matches the entire data set for all initial cell densities (Jin et al., 2016). One way of interpreting this result is that the cells in the scratch assay do not proliferate logistically.

In the present work, we use a combined experimental and mathematical approach to investigate whether the proliferation of cells in a scratch assay can be modelled with the classical logistic equation. Our approach involves performing a series of proliferation assays to act as a control so that we can examine whether the process of scratching the monolayer affects the way that cells proliferate. While many experimental studies implicitly assume that scratching the monolayer does not affect cell proliferation, others suggest the process of scratching can trigger certain signalling pathways that may have some effects on the way that cells proliferate (Nikolić et al., 2006;Nishio et al., 2005). To investigate these questions, we perform a suite of scratch assays and proliferation assays using the IncuCyte ZOOM™ system (Johnston et al., 2015). For both types of assays, we use the PC–3 prostate cancer cell line (Kaighn et al., 1979), and we consider varying the initial seeding condition so that we can examine the influence of varying the initial cell density.

To quantitatively test the suitability of the logistic growth model, we extract cell density information from the experimental images and then estimate the per capita growth rates from the data for both the scratch assays and the proliferation assays. Our results show that the evolution in cell density in the proliferation assays appears to be logistic for the entire duration of the experiment. In contrast, the variation in cell density in the scratch assays is very different. We observe two phases in the scratch assays: (i) a disturbance phase at early time, in which the proliferation of cells is not logistic; and, (ii) a classic logistic growth phase for the remainder of the experiment. These two phases are observed in all of our experiments, across a wide range of initial cell densities. The differences how cells proliferate in the scratch assay and the proliferation assay is surprising because we are making observations well away from the location of the scratch. This finding that we have two phases of proliferation in scratch assays is significant because many mathematical studies implicitly assume that cells in scratch assays proliferate logistically for the entire duration of the experiment (Maini et al., 2004a; Maini et al., 2004b; Savla et al., 2004; Cai et al., 2007;Sengers et al., 2007; Tremel et al., 2009; Johnston et al., 2015; Jin et al., 2016). However, our finding is that cells located far away from the scratch proliferate very differently to cells in the proliferation assay.

This manuscript is organised in the following way. First, we describe the experimental methods, including how we process the experimental images to obtain cell density information. We then outline the logistic growth model and the least–squares method for calibrating the model to our data. By presenting information about the evolution of the cell density and the per capita growth rate, we identify two phases of proliferation in the scratch assays. These phases are identified by focusing on regions of the scratch assay that are located well behind the location of the scratch. After calibrating the solution of the logistic model to the cell density information, our results suggest that the logistic equation is relevant for the proliferation assays but only for the later phase in the scratch assays. We conclude this study by discussing some of the limitations, and we outline some extensions for future work.

## 2 Methods

### 2.1 Experimental Methods

We perform scratch assays and proliferation assays using the IncuCyte ZOOM™ live cell imaging system (Essen BioScience, MI USA). All experiments are performed using the PC–3 prostate cancer cell line (Kaighn et al., 1979). These cells, originally purchased from American Type Culture Collection (Manassas, VA, USA), are a gift from Lisa Chopin (April, 2016). The cell line is used according to the National Health and Medical Research Council (NHMRC) National statement on ethical conduct in human research with ethics approval for Queensland University of Technology Human Research Ethics Committee (QUT HREC 59644, Chopin). Cells are propagated in RPMI 1640 medium (Life Technologies, Australia) with 10% foetal calf serum (Sigma–Aldrich, Australia), 100 U/mL penicillin, and 100 /g/mL streptomycin (Life Technologies), in plastic tissue culture flasks (Corning Life Sciences, Asia Pacific). Cells are cultured in 5% CO_2_ and 95% air in a Panasonic incubator (VWR International) at 37 ^o^C. Cells are regularly screened for *Mycoplasma* (Nested PCR using primers from Sigma–Aldrich).

Cell counting is performed using a Neubauer–improved haemocytometer (ProSciTech, Australia). Cells, grown to approximately 80% confluence, are removed from the flask using TrypLE™ (Life Technologies) in phosphate buffered saline (pH 7.4) and resuspended in culture medium ensuring that they are thoroughly mixed. After resuspension, an aliquot of 10 μL is quickly removed before the cells start to settle. A 1:1 mixture of cell suspension and 0.4% trypan blue solution (Sigma–Aldrich; a blue stain that is only absorbed by dead cells) is prepared and 10 AL of the solution is loaded onto the counting chamber of a clean Neubauer–improved haemocytometer. The counting chamber of a haemocytometer is delineated by grid lines that identify four chamber areas to be used in cell counting. The volume of the chamber area is 1a10^4^ mL. Using a microscope, each chamber area is viewed, and the live cells that are not coloured in blue are counted. The cell density is calculated by taking the average of the four readings and multiplying it by 10^4^ and the dilution factor, to obtain the approximate number of cells per mL of the cell suspension (Louis and Siegel, 2011).

For the proliferation assays, the cell count is determined and the cells are seeded at various densities in 96–well ImageLock plates (Essen Bioscience). Cells are distributed in the wells of the tissue culture plate as uniformly as possible. We report results for initial seeding densities of approximately 12,000, 16,000 and 20,000 cells per well. After seeding, cells are grown overnight to allow for attachment and some subsequent growth. The plate is placed into the IncuCyte ZOOM™ apparatus, and images are recorded every two hours for a total duration of 48 hours. An example of a set of experimental images from a proliferation assay is shown in Figure 1a–c. For each initial seeding condition we perform 16 identically prepared experimental replicates (*n* = 16).

For the scratch assays, the cell count is determined and the cells are seeded at various densities in 96–well ImageLock plates (Essen Bioscience). Cells are distributed in the wells of the tissue culture plate as uniformly as possible. We report results for initial seeding densities of approximately 12,000, 16,000 and 20,000 cells per well. After seeding, cells are grown overnight to allow for attachment and some subsequent growth. We use a WoundMaker™ (Essen BioScience) to create uniform scratches in each well of a 96–well ImageLock plate. To ensure that all cells are removed from the scratched region, a modification is made to the manufacturer’s protocol, where the scratching motion is repeated 20 times over a short duration before lifting the WoundMaker™. After creating the scratch, the medium is aspirated and the wells are washed twice with fresh medium to remove any cells from the scratched area. Following the washes, 100 eL fresh medium is added to each well and the plate is placed into the IncuCyte ZOOM™ apparatus. Images of the collective cell spreading are recorded every two hours for a total duration of 48 hours. An example of a set of experimental images taken from a scratch assay is shown in Figure 1d–f. For each initial seeding condition we perform 16 identically prepared experiments in different wells of the tissue culture plate (*n* = 16). Throughout this work we will refer to these identically prepared experiments in different wells as different *replicates.*

**Fig 1.**
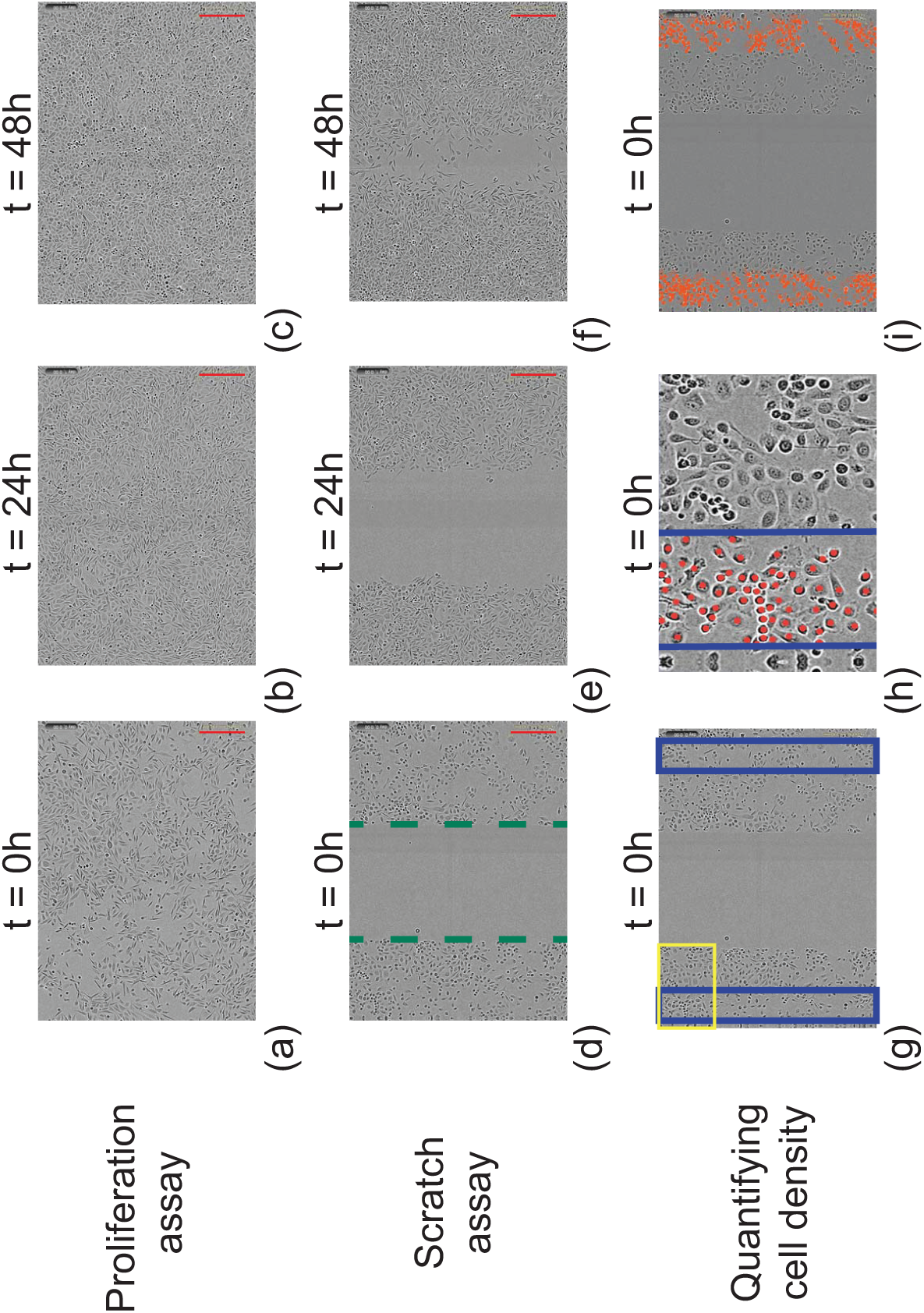
Experimental images. (a)–(c) A summary of IncuCyte ZOOMTM experiments for proliferation assays. (d)–(f) A summary of IncuCyte ZOOMTM experiments for scratch assays. Images show both types of experiments initiated with 16,000 cells per well. The time at which the image is recorded is indicated on each subfigure, and the scale bar (red line) corresponds to 300μm. The image in (d), at t = 0 hours, shows the approximate location of the position of the leading edges (dashed green). (g)–(i) To quantify the cell density profile, two rectangles of width 200*μ*m, are superimposed on the experimental image as shown in (g). Manual cell counting is used to estimate the number of cells in each subregion, and these estimates are converted into an estimate of cell density in these regions at two–hour intervals during the first 18 hours of the experiment, and then at six–hour intervals during the remaining 30 hours of the experiment. To count individual cells we zoom in to focus on certain subregions, such as shown in (h), which corresponds to the yellow rectangle highlighted in (g). Using the counting features in Adobe Photoshop (Adobe Systems Incorporated, 2016), we identify individual cells and place a unique marker on each cell (red disk), as shown in (h). After each image is processed in this way we have identified the total number of cells in the two subregions in the image, as shown in (i), and then we convert these estimates of cell numbers into an estimate of cell density.

### 2.2 Experimental Image Processing

To obtain cell density information from the experimental images, we count the number of cells in two identically sized subregions that are well behind the location of the scratch, as shown in Figure 1g. The positions of the two subregions are located about 400 tm behind the scratch, and each subregion has dimensions 1430 mm x 200 mm. Throughout this work, we refer to the subregion to the left of the image as subregion 1, and the subregion to the right of the image as subregion 2. Because the subregions are located well away from the scratched region, we are able to invoke a simplifying assumption that the dynamic changes in cell density in these subregions is due to cell proliferation alone (Supplementary Material) (Johnston et al., 2015). We do not use data that are directly adjacent to the left or right sides of the images since this corresponds to the boundary of the field of view. Cells in each subregion are counted in Photoshop using the ‘Count Tool’ (Adobe Systems Incorporated, 2016). After counting the number of cells in each subregion, we divide the total number of cells by the total area to give an estimate of the cell density. We repeat this process for each replicate and calculate the sample mean of the cell density at two–hour intervals during the first 18 hours of the experiment where the most rapid temporal changes take place. Then, during the last 30 hours of the experiment, we count cells at six–hour intervals.

One of the assumptions we make when analysing data from the scratch assay is that the two subregions are sufficiently far away from the edges of the scratch so that there are no spatial variations in cell density at these locations for the entire duration of the experiment. This assumption allows us to attribute any changes in cell density in the subregions to be a result of cell proliferation (Johnston et al., 2015). Quantitative evidence to support this assumption is provided in the Supplementary Material document.

### 2.3 Mathematical Methods

The logistic growth equation, given by Equation (1), has an exact solution

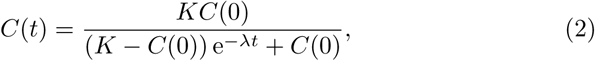

which is a sigmoid curve that monotonically increases from the initial density *C*(0) to *K* as *t* →∞. An important feature of the logistic growth model that we will make use of in this study is that the per capita growth rate, (1/*C*)(d*C*/d*t*) = λ(1 – *C*/K), decreases linearly with *C*.

We estimate the two parameters in the logistic growth model, λ and *K*, by minimising a least–squares measure of the discrepancy between the solution of the logistic growth equation and the average cell density information in our subregions that are located far away from the scratched region. The least– squares error is given by

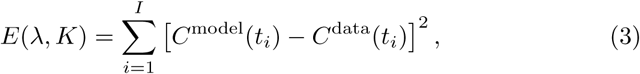

where *i* is an index that indicates the number of time points used from the experimental data sets and *I* is the total number of time points used in the calibration procedure. We calibrate the solution of the logistic growth equation to the average cell density information using the MATLAB function lsqcurvefit (MathWorks, 2016) that is based on the Levenberg–Marquardt algorithm. For notational simplicity we denote the minimum least–squares error as *E*_min_ = E (λˉ *K*ˉ). Each time we use the MATLAB function lsqcurvefit, we always check that the least-squares estimates of λˉ and *K*ˉ are independent of the initial estimate that is required for the iterative algorithm to converge.

### 3 Results and Discussion

#### 3.1 Quantitative assessment of experiments

##### 3.1.1 Initial cell density

Many previous studies that calibrate solutions of mathematical models to experimental data from proliferation or scratch assays make use of just one initial density of cells (Cai et al., 2007; Tremel et al., 2009; Maini et al., 2004a; Maini et al., 2004b). To provide a more thorough investigation of the suitability of various mathematical models, we calibrate mathematical models to a suite of experimental data where the initial density of cells is intentionally varied (Jin et al., 2016). To achieve this, our experimental procedure involves placing a different number of cells into each well of the tissue culture plate. We describe this as varying the *initial seeding condition.* In this work we consider three different initial seeding conditions that correspond to placing either: (i) 12,000; (ii) 16,000; or, (iii) 20,000 cells per well. For brevity, we refer to these three conditions as initial seeding conditions 1, 2 and 3, respectively.

After a particular number of cells are placed into the tissue culture plate, the cells are incubated overnight to allow them to attach to the plate and begin to move and proliferate. The experiments are then performed on the following day. Since the cell density changes overnight, we will refer to the initial density of cells at the beginning of the experiment on the following day, as the *initial cell density.* Intuitively, we expect that the initial cell density in proliferation assays will be greater than the cell density associated with the initial seeding condition, because the cells have had a period of time to attach and begin to proliferate.

Before we examine the temporal evolution of cell density in our experiments, we first examine the variability in the initial cell densities amongst our various experimental replicates. This is essential, since the process of placing either 12,000, 16,000 or 20,000 cells in each well of the tissue culture place is, at best, an approximation. To quantify the variability in the initial cell density, we count the number of cells in the two subregions, as shown in Figure 1g, and convert these counts into an estimate of the initial cell density, *C*(0). We repeat this procedure for both the proliferation and scratch assays, giving a total of 96 individual estimates of the initial cell density. These 96 estimates of the initial cell density are reported in Figure 2, revealing three features:

1. In general, those experiments initiated with a higher number of cells per well lead to a higher initial cell density after the overnight attachment and proliferation has taken place;
2. Within each initial seeding condition, the variability in initial cell density for the proliferation assays is very similar to the variability in initial cell density for the scratch assays; and,
3. There is a large variation in the initial cell density within each initial seeding condition.

Of these three features, the variation in the initial cell density within each initial seeding condition is very important. For example, the greatest recorded initial cell density for initial seeding condition 1 (12,000 cells per well) is greater than the smallest recorded initial cell density for initial seeding condition 3 (20,000 cells per well). This means that we ought to take great care when selecting particular experimental replicates from the 96 data sets in Figure 2, otherwise our results could be misleading when we try to examine how the results depend on the initial cell density.

**Fig 2.**
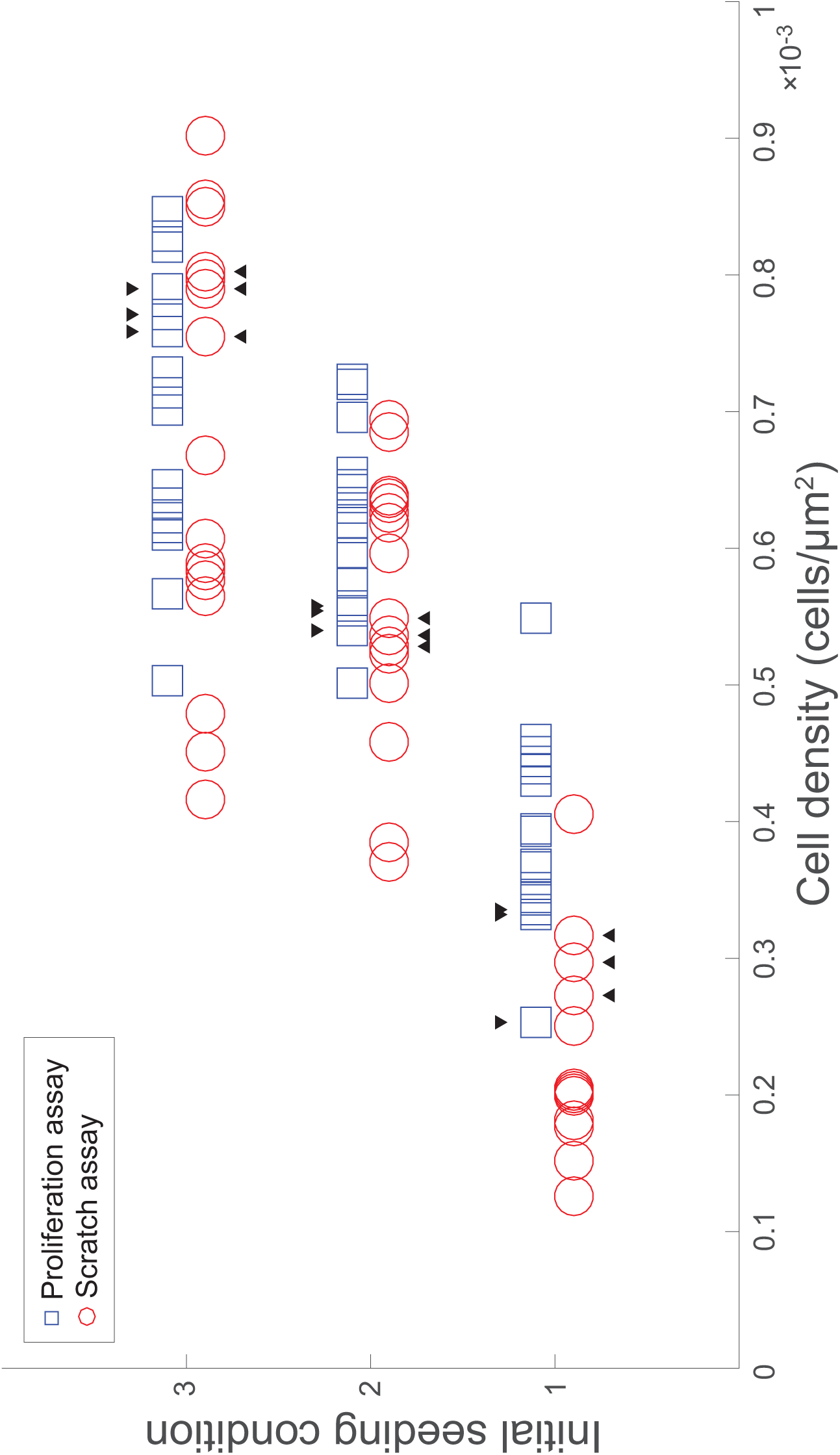
Variation of initial cell densities. Initial cell densities in the 96 replicates of proliferation and scratch assays. Results for the three different initial seeding conditions are shown. Initial seeding condition 1 corresponds to 12,000 cells per well; initial seeding condition 2 corresponds to 16,000 cell per well; and initial seeding condition 3 corresponds to 20,000 cells per well. For each initial seeding condition, each experiment is repeated n = 16 times and the variation in initial cell density is illustrated by comparing the spread of estimates of cell density on the horizontal axis. Each blue square represents an individual replicate of the proliferation assay, and each red circle represents an individual replicate of the scratch assay. The 18 black triangles indicate the individual replicates chosen to construct the cell density information.

We select three replicates from each initial seeding condition for both the proliferation and scratch assays so that the initial cell density for the initial seeding condition 3 is greater than the initial cell density for the initial seeding condition 2, which is greater than the initial cell density for the initial seeding condition 1. Furthermore, we select three replicates for both the proliferation and scratch assays from each initial seeding condition. These choices are made so that the initial cell density for each type of assay is approximately the same within each seeding condition. To satisfy these constraints we choose three replicates from each set of 16 experimental replicates. The selected replicates are indicated in Figure 2.

##### 3.1.2 Cell density information

Using the previously identified three experimental replicates for each type of assay and each initial seeding condition (Figure 2), we plot the evolution of the cell density as a function of time for each experimental replicate, as shown in Figure 3. We also superimpose, in Figure 3, the evolution of the average cell density for each type of assay and each initial seeding condition. We see that the differences in initial density between the proliferation assay and the scratch assay are minimal. The most obvious trend in the data is that the cell density in both the proliferation assay and the scratch assay increases dramatically with time, regardless of the initial condition. It is worth emphasizing that seeding condition 1 involves a relatively small initial cell density, whereas seeding conditions 2 and 3 are not particularly small. For example, the initial cell density for initial seeding condition 3 is approximately equal to the cell density for initial seeding condition 1 after a period of 24 hours has elapsed. Therefore our experimental design allows us to make a clear distinction between the effects of small cell density, which would appear more strongly and for longer in initial seeding condition 1 than initial seeding condition 3, and the effects of early time, which would appear equally in all three initial seeding conditions. We return to this issue later.

**Fig 3.**
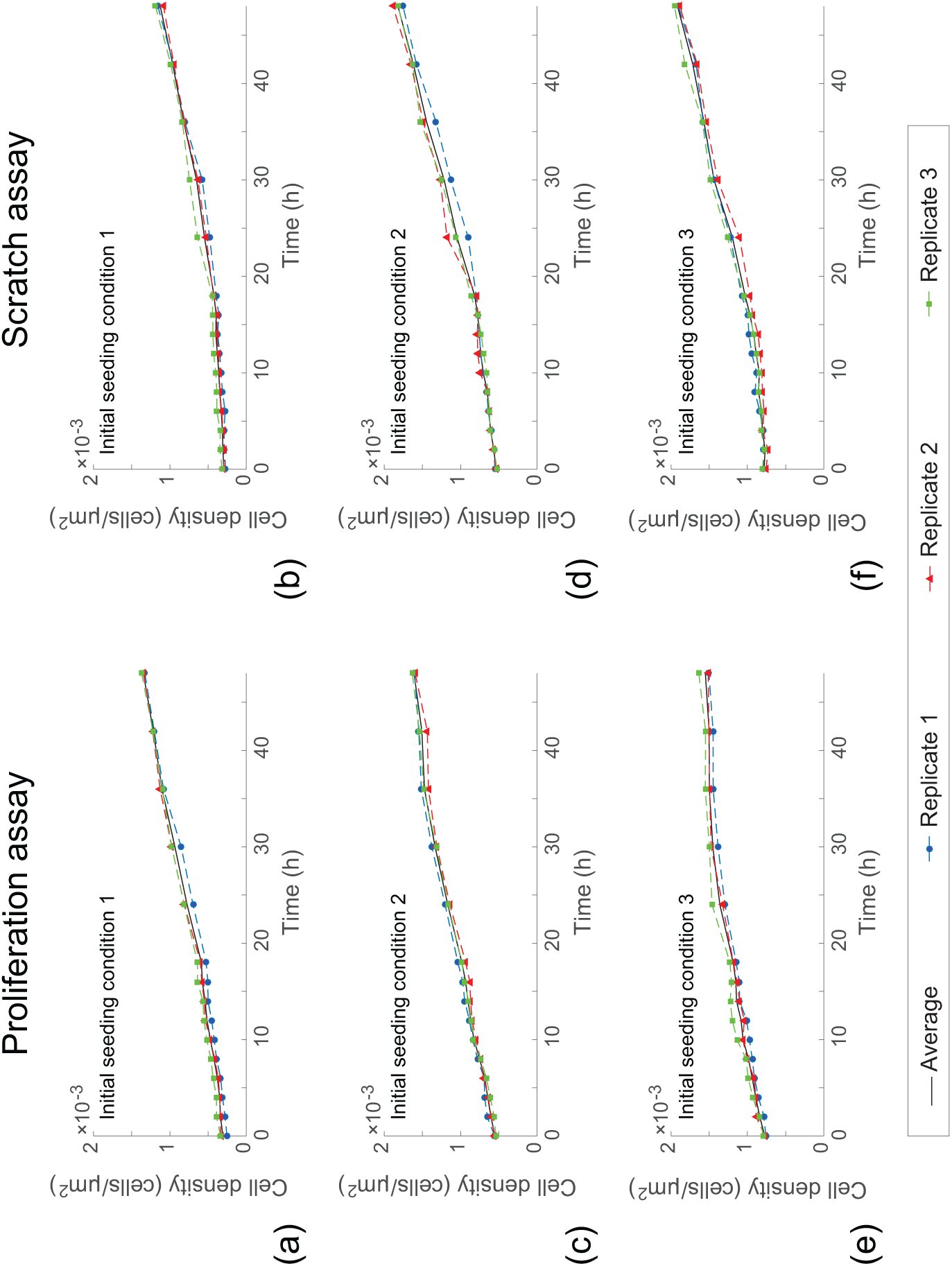
Temporal evolution of cell density. Results in (a)–(f) correspond to proliferation and scratch assays initiated with 12,000 (initial seeding condition 1); 16,000 (initial seeding condition 2); and, 20,000 (initial seeding condition 3) cells per well, as indicated. Cell density profiles are shown at two–hour intervals during the first 18 hours, and at six–hour intervals during the remaining 30 hours of the experiment. For each experiment, we report results for three identically prepared experimental replicates, and the average of these three data sets is also shown.

We note that it could be possible to calibrate the solution of Equation (1) to any of the density curves in Figure 3, and this approach has been widely used (Cai et al., 2007; Tremel et al., 2009; Simpson et al., 2013; Treloar and Simpson, 2013). However, there is no guarantee that simply fitting the solution of the logistic equation to this kind of data means that the logistic model describes the underlying mechanism (Simpson et al., 2014). To provide further insight into whether the logistic model applies to these data, we re-interpret the data in terms of the per capita growth rate.

#### 3.2 Per capita growth rate

To estimate the per capita growth rate, (1/*C*)(d*C*/d*t*), we use the cell density data in Figure 3 to estimate *dC/dt* using a finite difference approximation. Our estimate of d*C*/d*t* at the first and last time points is obtained using a forward and backward difference approximation, respectively, while our estimates at all other time points are obtained using an appropriate central difference approximation (Chapra and Canale, 2010). With these estimates, we plot the per capita growth rate as a function of the density in Figure 4. Results are shown for both proliferation and scratch assays, for the three initial densities considered.

**Fig 4.**
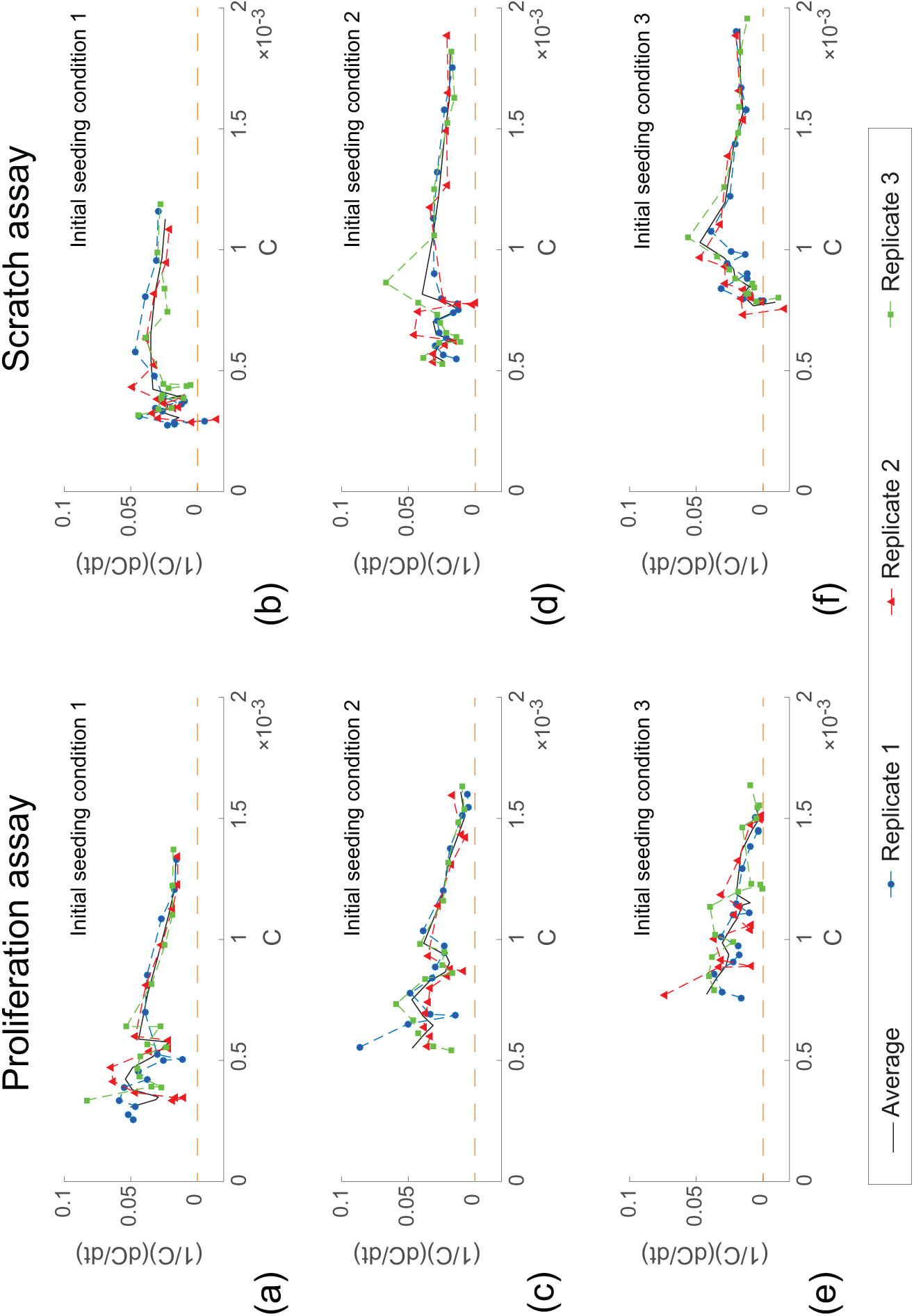
Per capita growth rates as a function of cell density. Results in (a)–(f) correspond to proliferation and scratch assays initiated with 12,000 (initial seeding condition 1); 16,000 (initial seeding condition 2); and, 20,000 (initial seeding condition 3) cells per well, as indicated. Per capita growth rate data is calculated using the data in Figure 3. For each experiment, we report results for three identically prepared experimental replicates, and the average of these three experimental replicates is also shown.

To interpret our results, it is instructive to recall that the data in Figure 3 show that the cell density, in each type of experiment for all three initial densities of cells, increases with time. Therefore, when we interpret each plot showing the per capita growth rate as a function of density in Figure 4, it is useful to recall how the data in these plots vary with time during the experiment. Data for smaller values of *C* in each subfigure in Figure 4 correspond to the early part of the experiment, and hence small *t*. In contrast, data for larger values of *C* in each subfigure in Figure 4 correspond to the latter part of the experiment, and hence larger *t*.

If the logistic growth model is valid, then we expect that the per capita growth rate will be a linearly decreasing function of the density. In contrast, other kinds of sigmoid-shaped carrying-capacity limited growth models, are associated with a a non-linear relationship between the per capita growth rate and the density. Visual inspection of the per capita growth rate data in Figure 4 reveals several trends:

1. The relationship between the per capita growth rate and the density in the proliferation assay is very different to the relationship between the per capita growth rate and the density in the scratch assay;
2. The relationship between the per capita growth rate for each proliferation assay, at each initial seeding condition, appears to be reasonably well approximated by a linearly decreasing function of density; and,
3. The relationship between the per capita growth rate for each scratch assay is more complicated, with the per capita growth rate increasing with density when the density is small, and then decreasing with density when the density is sufficiently large.

These observations suggest that the proliferation of cells in the scratch assay is very different to the proliferation of cells in a proliferation assay. Because we are examining the proliferation of cells that are located well away from the scratch, this result implies that the process of scratching the monolayer can induce non-local effects.

Instead of relying on visual interpretation alone, we now attempt to match the per capita growth rate data and the logistic growth model by fitting a series of straight lines to the averaged per capita growth rate data using lsqcurvefit (MathWorks, 2016). Results in Figure 5 show the least–squares straight line and the coefficient of determination for each data set. Results for the proliferation assay (Figure 5a, c and e) suggest that the putative linear relationship is reasonable since our straight lines have negative slope and the coefficient of determination is reasonably high (*R*^2^ = 0.50 – 0.87). In contrast, results for the scratch assay (Figure 5b, d and f) show that the least–squares linear regression is a poor match to the data with a very low coefficient of determination (*R*^2^ = 0.04 – 0.16). Indeed, the least–squares straight lines in Figure 5b and f are particularly troublesome since they have a positive slope which is biologically unrealistic, suggesting that the quantity λ/*K* is negative. Therefore, it is clear that the per capita growth data for the scratch assays does not follow a linearly decreasing straight line for the entire duration of the experiment, and the commonly-invoked logistic model does not appear to match these data at all.

**Fig 5.**
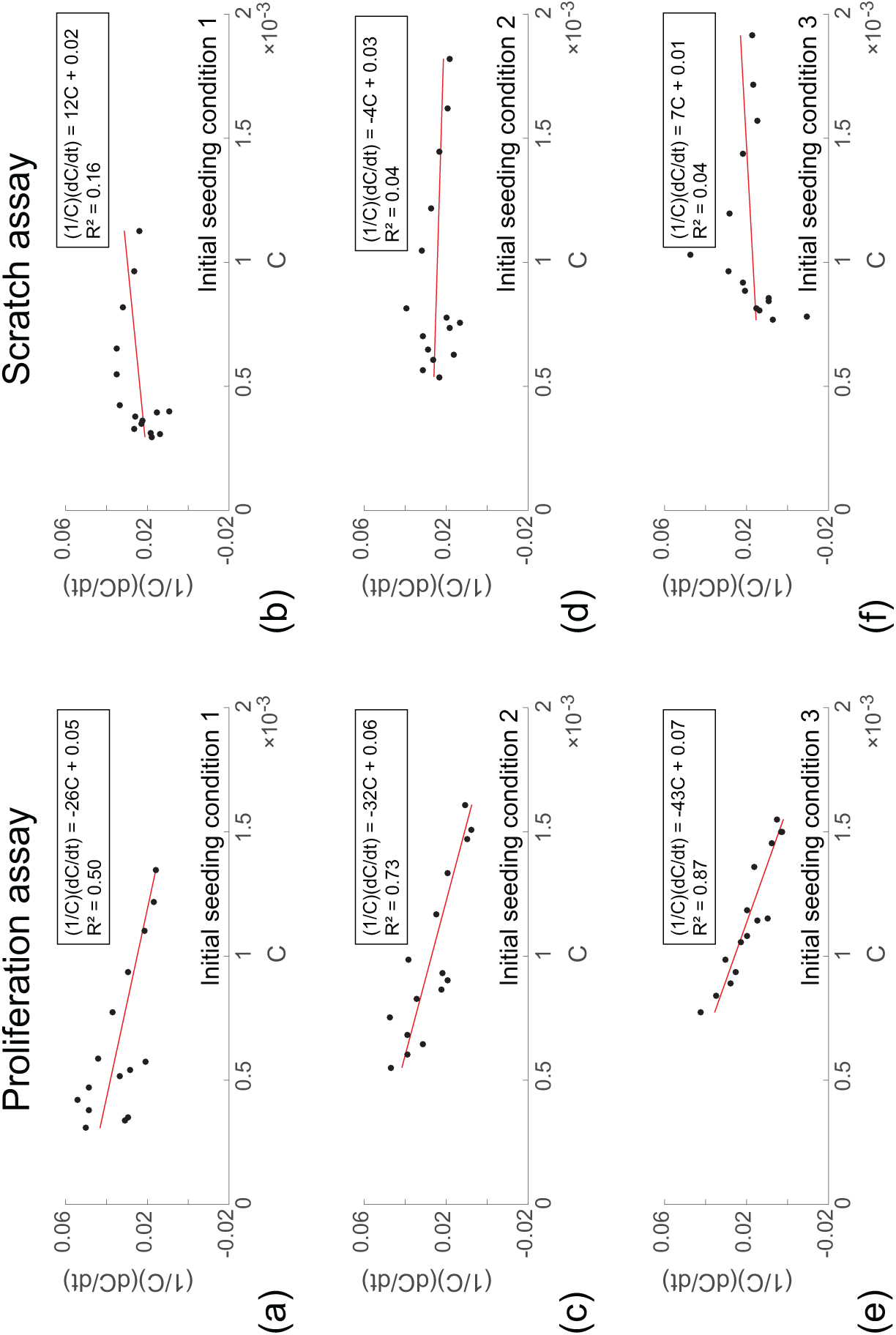
Least–squares straight line fit to per capita growth rate data for the entire duration of the experiment. Results in show the average per capita growth rate data as a function of density for both proliferation and scratch assays initiated with 12,000 (initial seeding condition 1); 16,000 (initial seeding condition 2); and, 20,000 (initial seeding condition 3) cells per well, as indicated. Black dots correspond to averaged data for the entire duration of the experiment. Solid lines show the least–squares linear relationship between the averaged per capita growth rate and averaged density, with *R*^2^ indicating the coefficient of determination.

The fact that we observe two very different trends in the per capita growth rate data for the scratch assay motivates us to conjecture that the proliferation of cells in the scratch assay, far away from the location of the scratch, takes place in two phases. The first phase, which occurs at early time, involves the per capita growth rate increasing with density. This trend is the opposite of what we expect if the logistic growth model is valid and not what we observe in the proliferation assay. The second phase, which occurs at later time, involves the per capita growth rate decreasing with the density. These two phases occur consistently across all three initial seeding conditions (Figure 4b, d and f). A schematic illustration of the differences observed between the per capita growth rate in the scratch assay and the proliferation assay is given in Figure 6. In this schematic, we refer to the first phase in the scratch assay as the *disturbance phase,* and the second phase in the scratch assay as the *growth phase.* The per capita growth data in the proliferation assay appear to be similar to the growth phase of the scratch assay for the entire duration of the experiment.

**Fig 6.**
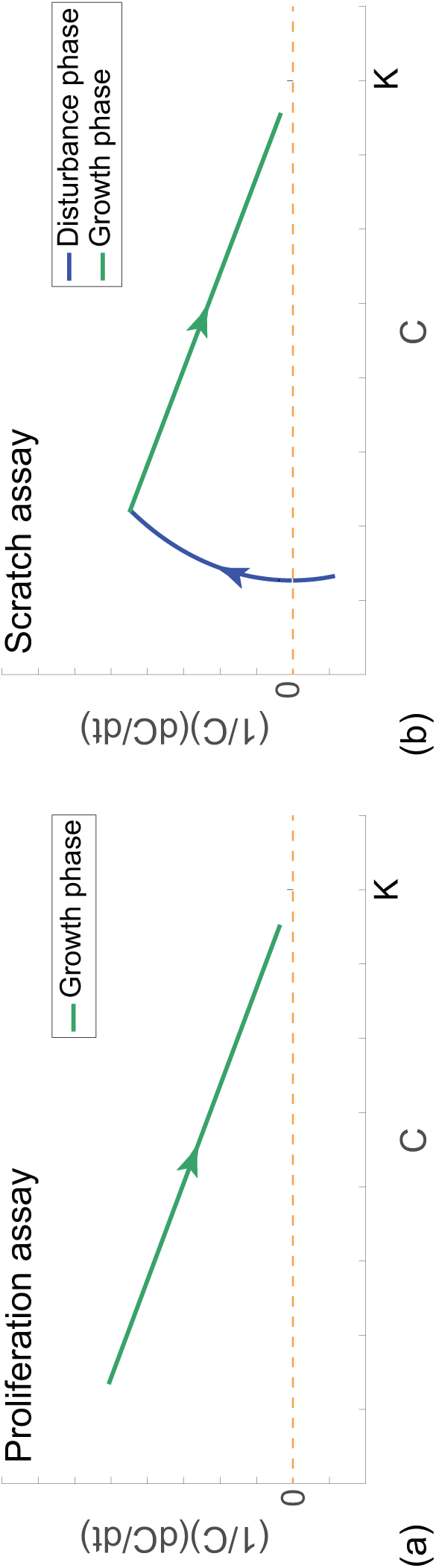
Schematic illustration of the differences between the proliferation and scratch assays. (a) Schematic showing the per capita growth rate as a function of density for the proliferation assays, (b) Schematic of the per capita growth rate for the scratch assays illustrating two phases of proliferation. The solid blue line indicates the disturbance phase in the scratch assay, and the solid green line indicates the growth phase in both the proliferation and scratch assays. The arrow heads indicate the direction of increasing time.

In the schematic (Figure 6), we suggest that the relationship between the per capita growth rate and the density during the growth phase is a linearly decreasing function, which is consistent with the logistic model. To quantitatively examine whether this assumption is valid for our data set we now construct a series of least–squares straight lines to our averaged per capita growth rate data during the growth phase. To examine this question, we need to quantitatively distinguish between the end of the first phase and the beginning of the second phase. We separate the data in Figure 5b, d and f into two groups, the disturbance phase for *t* < 18 hours, and the growth phase for *t* > 18 18 hours. To examine whether the data in the growth phase appear to be logistic we determine the least–squares linear relationship using lsqcurvefit (MathWorks, 2016) for the data in the growth phase. This least–squares straight line is superimposed on the averaged data for *t* ≥ 18 hours in Figure 7b, d and f. Again, a visual comparison of the match between the linear regression and the data in the growth phase, and the much higher values of coefficient of determination (*R*^2^ = 0.75 – 0.91) suggest that the putative linear relationship is reasonable.

**Fig 7.**
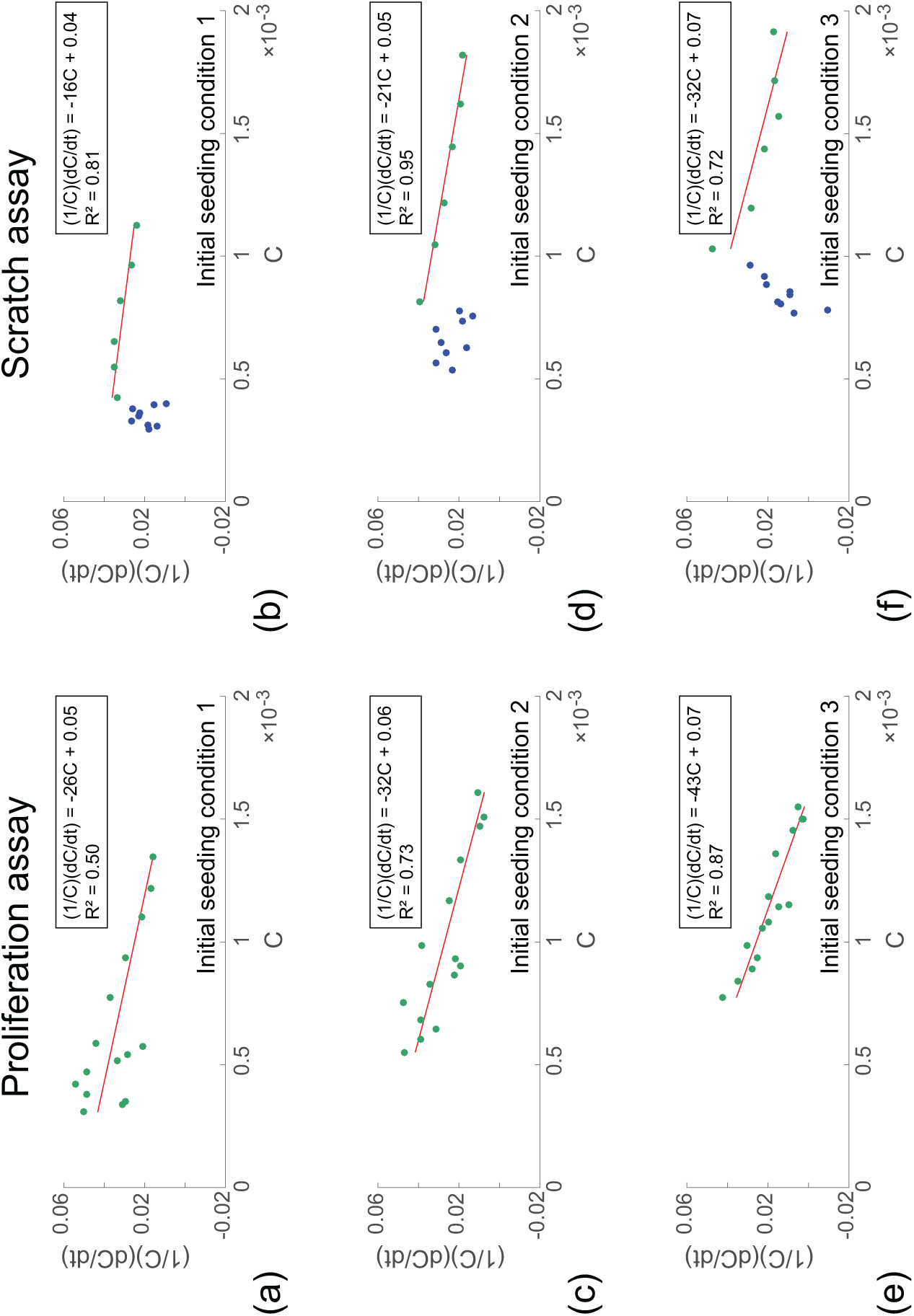
Straight line fit to per capita growth rates in the growth phase. Results in (a)–(f) show the average per capita growth rate data as a function of density for both proliferation and scratch assays initiated with 12,000 (initial seeding condition 1); 16,000 (initial seeding condition 2); and, 20,000 (initial seeding condition 3) cells per well, as indicated. Green dots correspond to averaged data in the growth phase (Figure 5), and blue dots correspond to averaged data in the disturbance phase (Figure 5). The solid lines show the least–squares linear relationship between the averaged per capita growth rate and averaged density, with *R^2^* indicating the coefficient of determination. The least–squares straight line is constructed using data from 0–48 hours in the proliferation assays, and using data from 18–48 hours in the scratch assays.

In summary, we have used the per capita growth rate information in Figure 4 to make a distinction between the disturbance phase and the growth phase in the scratch assay. These differences are highlighted in the schematic in Figure 6. Furthermore, guided by the observed relationship between the per capita growth rate and the density in the proliferation assay we assume that the logistic growth model applies and fit a straight line to the per capita growth rate data and find that the match between the data and the straight line appears to be reasonable. Similarly, we assume that the logistic growth model applies to the growth phase in the scratch assay, for *t* ≥ 18 hours. Fitting a straight line to the per capita growth data suggests that the logistic growth model is reasonable in the growth phase for the scratch assay. Now that we have used the per capita growth rate data to identify the disturbance and growth phases in the scratch assay, as well as providing evidence that cells proliferate logistically in the growth phase, we re–examine the cell density profiles with a view to estimating λ and *K*.

#### 3.3 The logistic growth model

To calibrate the logistic growth model to our data from the proliferation assay, we match the solution of Equation (1) to the averaged data in Figure 3a, c and over the entire duration of the experiment, 0 ≤ *t* ≤ 48 hours. To calibrate the logistic growth model to our data from the scratch assay, accounting for the differences in the disturbance and growth phases, we match the solution of Equation (1) to the averaged data in Figure 3b, d and f during the growth phase only, 18 ≤ *t* ≤ 48 hours. This provides us with six estimates of λ̅ and *K̄*. To demonstrate the quality of the match between the experimental data and the calibrated logistic model, we superimpose the experimental data and Equation (2) with λ̅= *K̅* and *K̅* = *K*, for each initial seeding condition and for both assays in Figure 8. These results show that the quality of match between the solution of the calibrated model and the experimental data is excellent. Our estimates of λ and *K̅* are summarised in Tables 1 and 2 for the proliferation assay and the scratch assay, respectively. In summary, our estimates of λ vary within the range λ = 0.048 – 0.067 h^−1^, and our estimates of *K* vary within the range *K* = 1.6 – 2.5 × 10^−3^ cells//m^2^. Strictly speaking, since λ and *K* are supposed to be constants in Equation (1), the fact that we see only a relatively small variation in our estimates of these parameters is encouraging. In particular, we also report, in Tables 1 and 2, the sample standard deviation showing the variability of our estimates. Overall, we find that the coefficient of variation is approximately 10%, which is relatively small when dealing with this kind of biological data (Vo et al., 2015).

**Fig 8.**
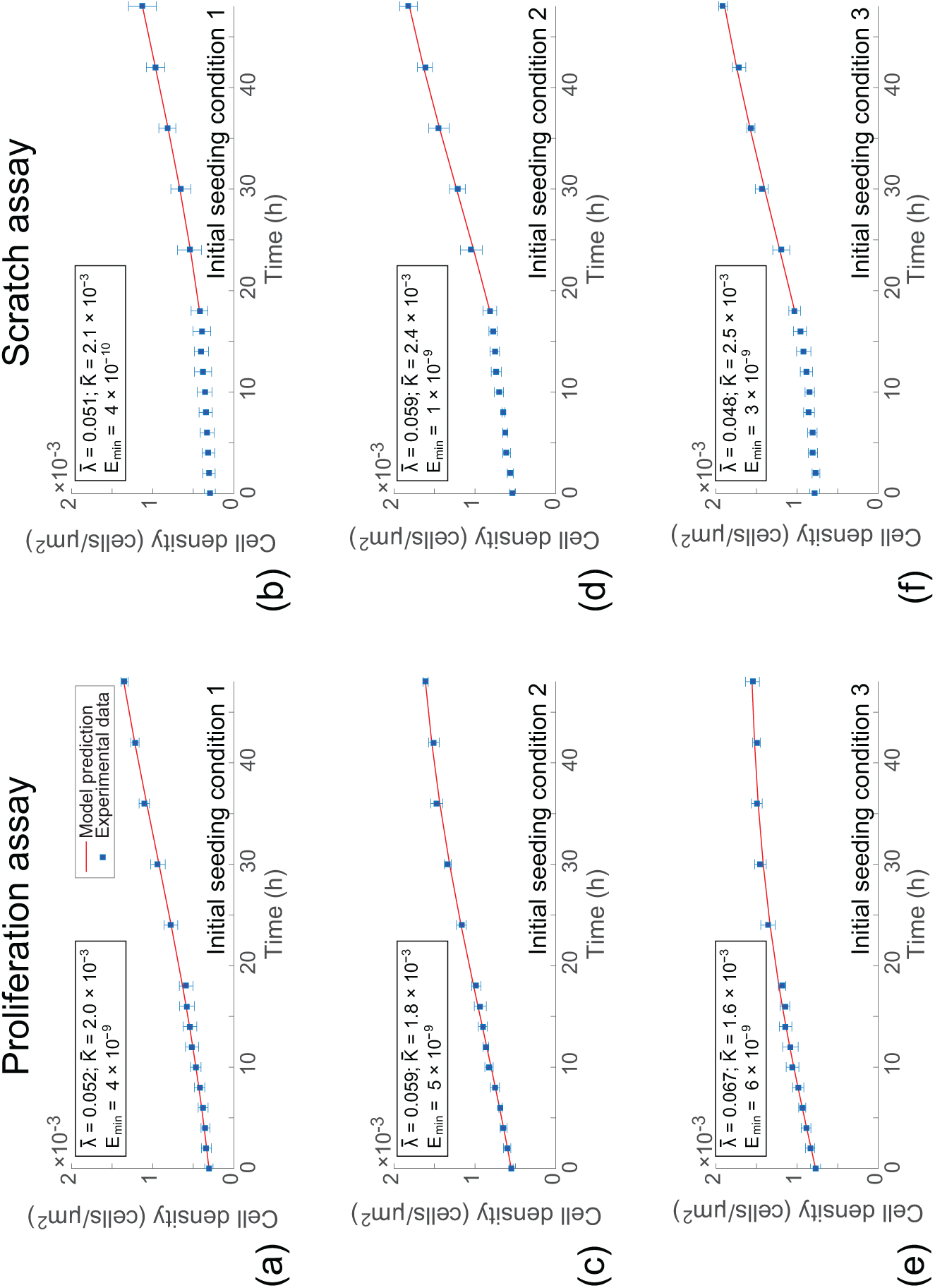
Calibrated solutions of the logistic growth equation using data from the growth phase. Results in (a)–(f) correspond to proliferation and scratch assays initiated with 12,000 (initial seeding condition 1); 16,000 (initial seeding condition 2); and, 20,000 (initial seeding condition 3) cells per well, as indicated. For each type of experiment the calibrated solution of the logistic growth equation (solid line) is compared to the experimental data in the growth phase (18–48 hours for scratch assays and 0–48 hours for proliferation assays). The least–squares estimates of λ and *K* are shown.

**Table 1.** Estimates of λ̅ and *K̅* for the proliferation assay using data from 0 < *t* 48 hours. All parameter estimates are given to two significant figures. Results are reported as the sample mean and the uncertainty is quantified in terms of the sample standard deviation.

| Initial seeding condition | $\bar{\lambda}$ (/h) | $\bar{K}$ (cells/$\mu\text{m}^2$) |
| --- | --- | --- |
| 1 | $0.052 \pm 0.004$ | $2.0 \times 10^{-3} \pm 8 \times 10^{-5}$ |
| 2 | $0.059 \pm 0.006$ | $1.8 \times 10^{-3} \pm 6 \times 10^{-5}$ |
| 3 | $0.067 \pm 0.009$ | $1.6 \times 10^{-3} \pm 2 \times 10^{-5}$ |
| Average | $0.059 \pm 0.008$ | $1.8 \times 10^{-3} \pm 2 \times 10^{-4}$ |

**Table 2.** Estimates of λ̅ and *K̅* for the scratch assay using data from 18 a *t* 48 hours. All parameter estimates are given to two significant figures. Results are reported as the sample mean and the uncertainty is quantified in terms of the sample standard deviation.

| Initial seeding condition | $\bar{\lambda}$ (/h) | $\bar{K}$ (cells/$\mu\text{m}^2$) |
| --- | --- | --- |
| 1 | $0.051 \pm 0.009$ | $2.1 \times 10^{-3} \pm 2 \times 10^{-3}$ |
| 2 | $0.059 \pm 0.02$ | $2.4 \times 10^{-3} \pm 1 \times 10^{-3}$ |
| 3 | $0.048 \pm 0.008$ | $2.5 \times 10^{-3} \pm 2 \times 10^{-4}$ |
| Average | $0.053 \pm 0.005$ | $2.3 \times 10^{-3} \pm 2 \times 10^{-4}$ |

We now explore how our estimates of λ and *K* are sensitive to whether or not we account for the differences in the disturbance and growth phases in the scratch assay. We repeat the same calibration process as described for the results in Figure 8, except now we take the standard, naive approach and calibrate the solution of Equation (1) to the averaged data in Figure 3b, d and f over the entire duration of the scratch assay, 0 ≤ *t* ≤ 48 hours. This procedure provides us with three additional estimates of λ̅ and *K̅* for the scratch assay, as summarised in Table 3.

**Table 3.** Estimates of λ̅ and *K̅* for the scratch assay using data from 0 ≤ *t* ≤ 48 hours. All parameter estimates are given to two significant figures. Results are reported as the sample mean and the uncertainty is quantified in terms of the sample standard deviation.

| Initial seeding condition | $\bar{\lambda}$ (/h) | $\bar{K}$ (cells/$\mu\text{m}^2$) |
| --- | --- | --- |
| 1 | $0.028 \pm 0.001$ | $2.8 \times 10^7 \pm 1 \times 10^7$ |
| 2 | $0.029 \pm 0.005$ | $8.7 \times 10^{-3} \pm 3 \times 10^6$ |
| 3 | $0.019 \pm 0.0002$ | $1.6 \times 10^7 \pm 6 \times 10^6$ |
| Average | $0.025 \pm 0.006$ | $1.5 \times 10^7 \pm 1 \times 10^7$ |

To demonstrate the quality of the match between the experimental data and the calibrated logistic model, we superimpose the experimental data and Equation (2) with λ = λ̅ and *K* = *K̅*, for each initial seeding condition and for both assays in Figure 9. When we visually compare the quality of the match between the experimental data in Figure 8 and Figure 9, and the corresponding calibrated solution of the logistic equation, there does not appear to be any significant difference at all. It is worth noting that the values of *E*_min_ in Figure 9b, d, and f are an order of magnitude greater than the corresponding values in Figure 8b, d and f. This implies that the match between the logistic model and the experimental data is improved when we ignore that data during the disturbance phase. However, at first glance, these differences are visually indistinguishable when we compare the results in Figure 8 and 9. In contrast, when we examine the estimates of *K̅* and λ̅ in Table 3, the importance of properly accounting for the disturbance phase in the scratch assay becomes strikingly obvious. For example, taking this latter approach, our estimates of the carrying capacity vary within the range K = 1.6 × 10^−3^ – 2.8 10^7^ cells//m^2^, and our estimates of the proliferation rate vary within the range λ = 0.019 – 0.067 h^−1^. We recall that λ and *K* are supposed to be constants in Equation (1), and so the fact that this naive calibration process suggests that the least–squares estimate of the carrying capacity density varies of many order of magnitude provides a clear illustration that this standard approach to calibrating the logistic equation to our experimental data is problematic. We note that the results of the Levenberg–Marquardt algorithm are robust, returning the same least-squares estimates of λ̅ and *K̅* for any positive initial estimate of *K* and λ in the iterative algorithm (MathWorks, 2016).

**Fig 9.**
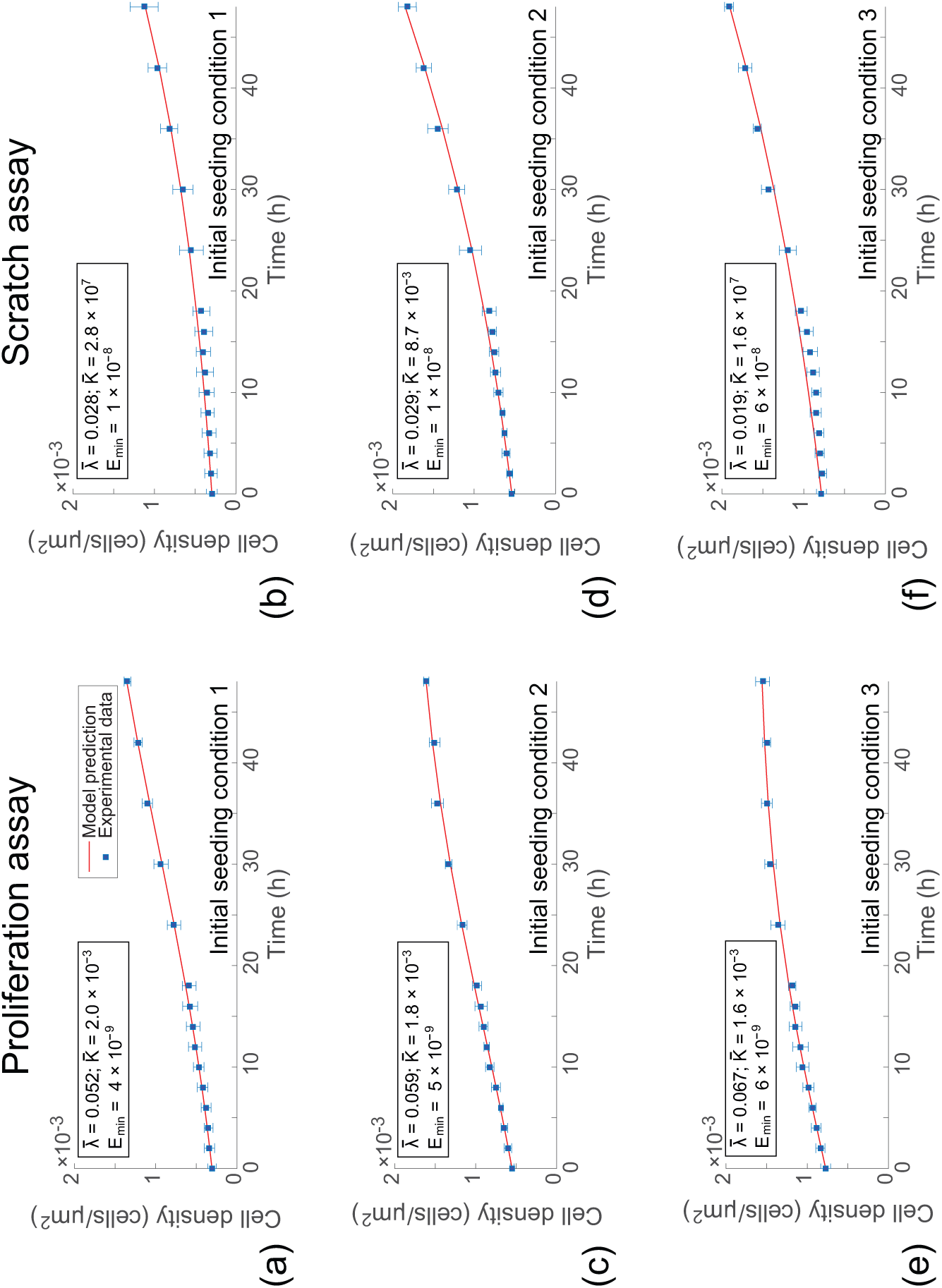
Calibrated solutions of the logistic growth equation using the entire data set. Results in (a)–(f) correspond to proliferation and scratch assays initiated with 12,000 (initial seeding condition 1); 16,000 (initial seeding condition 2); and, 20,000 (initial seeding condition 3) cells per well, as indicated. For each type of experiment the calibrated solution of the logistic growth equation (solid line) is compared to the entire experimental data set. The least–squares estimates of λ̅ and *K̅* are shown.

Comparing the ranges of estimates for λ and *K* in Tables 2 and 3 shows that the model calibration procedure is extremely sensitive. For example, our range of estimates of *K* when we account for the disturbance phase is smaller than a factor of two amongst the six estimates. In contrast, when we neglect the disturbance phase, our estimates of *K* vary across more than ten orders of magnitude amongst the six estimates. Similarly, our range of estimates of λ when we account for the disturbance phase is smaller than a factor of 1.5 among the six estimates. Again, in contrast, when we take a standard approach and neglect the disturbance phase our estimates of λ vary by more than a factor of three amongst the six estimates.

## 4 Conclusion

In this work we investigate the suitability of the logistic growth model to describe the proliferation of cells in scratch assays. Scratch assays are routinely used to study the ability of a population of cells to re–colonise an initially vacant region on a two–dimensional substrate (Liang et al., 2007; Tremel et al., 2009; Kramer et al., 2013; Treloar and Simpson, 2013). Most experimental interpretations of scratch assays are made using relatively straightforward measurements (Liang et al., 2007). However, to provide additional insights into the mechanisms involved in the re–colonisation process, some previous studies have calibrated the solution of a reaction-diffusion equation to data from a scratch assay (Sheardown and Cheng, 1996; Maini et al., 2004a; Maini et al., 2004b; Savla et al., 2004; Cai et al., 2007; Sengers et al., 2007; Shakeel et al., 2013; Simpson et al., 2007; Johnston et al., 2015; Jin et al., 2016). In these reaction–diffusion equations, it is commonly assumed that carrying capacity–limited proliferation of cells can be described by a logistic growth model. However, the suitability of this assumption is rarely examined beyond the process of simply calibrating the solution of the relevant model to match the experimental data.

To examine the suitability of the logistic growth model, we perform a series of scratch assays and proliferation assays for three different initial seeding densities. Cell proliferation assays are prepared in exactly the same way as a scratch assay, except that the monolayer of cells is not scratched. This allows us to treat the cell proliferation assays as a control experiment so that we can examine whether the process of artificially scratching the monolayer of cells affects the way that cells proliferate, even when those cells are located far away from the scratch. Instead of examining the dynamics of the cell density near the scratched region where there will be a net flux of cells into the vacant region (Jin et al., 2016), we quantify the cell density in two subregions that are located far behind the location of the scratch, where the cell density is approximately spatially uniform (Supplementary Material). This means that the temporal dynamics of the cell density in these subregions is due to cell proliferation only (Johnston et al., 2015).

We plot the time evolution of cell density, far away from the initially scratched region, in both the scratch and proliferation assays. To examine whether our results are sensitive to the initial density of cells, we repeat each experiment using three different initial cell densities. Plots of the evolution of the cell density are given over a total duration of 48 hours, and these plots appear to correspond to a series of sigmoid curves. At this point it would be possible to simply calibrate the solution of the logistic growth model to these data to provide an estimate of the proliferation rate, λ, and the carrying capacity density, *K*. This is a standard approach that has been used by us (Johnston et al., 2015) and many others (Cai et al., 2007; Sengers et al., 2007; Tremel et al., 2009). However, while this standard calibration procedure can be used to provide estimates of the parameters, this model calibration procedure does not provide any validation that logistic growth is relevant (Simpson et al., 2014).

Rather than calibrating the logistic growth model to our experimental data, we attempt to assess the suitability of the logistic growth model by converting the cell density evolution profiles into plots of the per capita growth rate as a function of density. We find that the plots of the per capita growth rate as a function of density reveal several key differences between the scratch and proliferation assays. If the logistic growth model is valid, then we expect to see a decreasing linear relationship between the per capita growth rate and the cell density for the entire duration of the experiment. While the plots of the per capita growth rate as a function of density for the proliferation assays appear to be consistent with the logistic model, the per capita growth rate data for the scratch assays are very different. For the scratch assay data, the per capita growth rate increases with cell density at low density during the early part of the experiment. This behaviour, which is observed for all three initial densities of cells in the scratch assays, is the opposite of what we would expect if the logistic growth model were valid. However, at higher cell densities during the latter part of the experiment, we observe that the per capita growth rate in the scratch assays appears to decrease, approximately linearly, with the cell density. This motivates us to propose that cell proliferation in a scratch assay involves two phases: (i) a *disturbance phase* in which proliferation does not follow the logistic growth model during the early part of the experiment; and, (ii) a *growth phase* where proliferation is approximately logistic during the latter part of the experiment. Guided by our per capita growth rate data, it appears that the disturbance phase in the scratch assays lasts for approximately 18 hours before the growth phase commences.

To estimate the parameters in the logistic growth model, we calibrate the solution of the model to our cell proliferation data for the entire duration of the experiment. This calibration procedure gives estimates of λ and *K* that are approximately consistent across the three initial conditions. We then calibrate the solution of the logistic growth model to the data from the growth phase in the scratch assay. This procedure also gives estimates of λ and *K* that are consistent across the three initial conditions, as well as being consistent with the estimates obtained from the cell proliferation assays. In contrast, if we take a naive approach and simply calibrate the solution of the logistic growth equation to the scratch assay data for the entire duration of the experiment, our estimates of λ and *K* vary wildly, despite the fact that the match between the experimental data and the calibrated solution of the logistic growth equation looks very good.

The results of our study strongly suggest that care ought to be taken when applying a logistic growth model, or a reaction-diffusion equation with a logistic source term, to describe scratch assays. Simply calibrating a mathematical model to experimental data might appear to produce an excellent match between the solution of the model and the experimental data, but this commonly-used procedure does not guarantee that the model is at all relevant (Simpson et al., 2014). Our results suggest that cell proliferation is impacted by the scratching procedure in a scratch assay, and that we require some time to pass before the disturbance phase ends. This is important because previous applications of logistic growth models and reaction-diffusion equations with logistic source terms have been calibrated to data from scratch assays without any regard for the disturbance phase (Cai et al., 2007; Tremel et al., 2009; Jin et al., 2016; Johnston et al., 2015).

It is also relevant to note that for the particular cell line we use, the disturbance phase that we identify lasts for approximately 18 hours. This is important because many scratch assays are performed for relatively short periods of time (Liang et al., 2007) and it is possible that standard experimental protocols do not allow for a sufficient amount of time to pass for the disturbance phase to end. Therefore, we suggest that scratch assays should be maintained for as long as possible so that sufficient time is allowed for the disturbance phase to pass.

It is worthwhile to note, and discuss, the fact that some of the features of our proposed two–phase growth model appear to be similar to the Allee effect (Allee and Bowen, 1932; Johnston et al., 2017; Lewis and Kareiva, 1993; Roques et al., 2012; Taylor and Hastings, 2005; Sewalt et al. 2016). Typically, Allee growth kinetics are normally invoked to describe some kind of low-density reduction in proliferation, relative to the logistic model (Lewis and Kareiva, 1993 Taylor and Hastings, 2005). The Allee growth model is given by

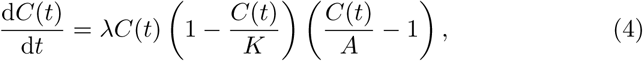

where the parameter *A* is called the Allee threshold. The key difference between the Allee growth model (Equation (4)) and the standard logistic model (Equation (1)), is the inclusion of the third factor on the right hand side of Equation (4). The incorporation of this factor has several consequences: (i) the growth rate is negative for *C*(*t*) < *A* (assuming *C*(*t*) < *K*); (ii) the growth rate is positive for *C*(*t*) > *A* (assuming *C*(*t*) < *K*); and, (iii) the relationship between the per capita growth rate and the density is quadratic. In many previous implementations of Allee growth models, an argument is made that the growth rate at small densities is reduced, relative to the logistic model, because of some kind of biological competition (Johnston et al., 2017;Lewis and Kareiva, 1993;Taylor and Hastings, 2005), corresponding to A ≪ K. Therefore, the Allee model is often used to represent reduced growth at small densities, *C*(*t*) ≪ K. The experimental data we present in Figures 4 and 7 are inconsistent with the Allee model for two reasons. First, the per capita growth data in Figure 4 corresponds to a reduced growth rate at early times during the experiment. This reduction in growth rate is observed across a range of initial densities, including seeing conditions 2 and 3 which do not involve small densities, as discussed in Section 3.1.2. Second, the per capita growth data in Figure 7 varies approximately linearly with density during the growth phase, whereas the Allee model implies that the relationship is quadratic.

One of the limitations of our study is that we have not identified the precise mechanism that causes the disturbance phase or the mathematical form of the disturbance phase. However it seems clear that the process of scratching a monolayer of cells has some impact on the proliferative behaviour of the cells away from the scratch, suggesting that either chemical or mechanical disturbance is transported throughout the experimental well as consequence of the scratching action. Regardless of the mechanism at play, our procedure of converting the cell density profiles into plots of the per capita growth rate allows us to identify the result of this disturbance. Another limitation of our work is that we deal only with one particular cell line, and it is not obvious how our estimate of the duration of the disturbance phase will translate to other cell lines. In this work we study the proliferation mechanism by converting the density data into per capita growth data and exploring whether the relationship between the per capita growth and density is approximately described by a linear function. We acknowledge that more sophisticated statistical techniques could be employed to provide further information (Dennis and Taper, 1994). However, since the main aim of this study is to explore the suitability of the logistic growth model for describing cell proliferation in a scratch assay, we do not pursue these more advanced statistical techniques here. Instead, we suggest that this could be the topic of a future study.

## 5 Acknowledgements

This work is supported by the Australian Research Council (DP140100249, FT130100148), and we appreciate the helpful comments from the two referees.

